# Aggression Subtypes Relate to Distinct Resting State Functional Connectivity in Disruptive Children and Adolescents

**DOI:** 10.1101/462382

**Authors:** Julia E Werhahn, Susanna Mohl, David Willinger, Lukasz Smigielski, Alexander Roth, Christoph Hofstetter, Philipp Stämpfli, Isabelle Häberling, Jilly Naaijen, Leandra M Mulder, Jeffrey C Glennon, Pieter J Hoekstra, Andrea Dietrich, Renee Kleine Deters, Pascal M Aggensteiner, Nathalie E Holz, Sarah Baumeister, Tobias Banaschewski, Melanie C Saam, Ulrike M E Schulze, David J Lythgoe, Arjun Sethi, Michael Craig, Mathilde Mastroianni, Ilyas Sagar-Ouriaghli, Paramala J Santosh, Mireia Rosa, Nuria Bargallo, Josefina Castro-Fornieles, Celso Aragno, Maria J Penzol, Barbara Franke, Marcel P Zwiers, Jan K Buitelaar, Susanne Walitza, Daniel Brandeis

**Affiliations:** Department of Child and Adolescent Psychiatry and Psychotherapy, University Hospital of Psychiatry Zurich, University of Zurich, Zurich, Switzerland; Neuroscience Center Zurich, University and ETH Zurich, Zurich, Switzerland; MR-Center of the Department of Psychiatry, Psychotherapy and Psychosomatics, University Hospital of Psychiatry Zurich, University of Zurich, Zurich, Switzerland; Radboud University Medical Center, Donders Institute for Brain, Cognition and Behavior, Department of Cognitive Neuroscience, Nijmegen, The Netherlands; Radboud University, Donders Institute for Brain, Cognition and Behavior, Centre for Cognitive Neuroimaging, Nijmegen, The Netherlands; University of Groningen, University Medical Center Groningen, Department of Child and Adolescent Psychiatry, Groningen, The Netherlands; Department of Child and Adolescent Psychiatry and Psychotherapy, Central Institute of Mental Health, Medical Faculty Mannheim/Heidelberg University, Mannheim, Germany; Department of Child and Adolescent Psychiatry/Psychotherapy, University Hospital, University of Ulm, Germany; Department of Neuroimaging, Institute of Psychiatry, Psychology & Neuroscience, King’s College London, London, UK; Department of Child Psychiatry, Institute of Psychiatry, Psychology & Neuroscience, King’s College London, London, UK; Child and Adolescent Psychiatry Department, Hospital Clinic of Barcelona, IDIBAPS, Barcelona, Spain; Clinic Image Diagnostic Center (CDIC), Hospital Clinic of Barcelona; Magnetic Resonance Image Core Facility, IDIBAPS, Barcelona, Spain; Child and Adolescent Psychiatry and Psychology Department, Institute Clinic of Neurosciences, Hospital Clinic of Barcelona, CIBERSAM, IDIBAPS, Department of Medicine, University of Barcelona, Barcelona, Spain; Child and Adolescent Psychiatry Department, Hospital General Universitario Gregorio Marañón School of Medicine, Universidad Complutense, IiSGM, CIBERSAM, Madrid, Spain; Radboud University Medical Center, Donders Institute for Brain, Cognition and Behavior, Department of Human Genetics, Nijmegen, The Netherlands; Radboud University Medical Center, Donders Institute for Brain, Cognition and Behavior, Department of Psychiatry, Nijmegen, The Netherlands; Karakter Child and Adolescent Psychiatry University Center, Nijmegen, The Netherlands; Zurich Center for Integrative Human Physiology, University of Zurich, Zurich, Switzerland

**Keywords:** reactive and proactive aggression, callous-unemotional traits, default mode network, amygdala, functional connectivity

## Abstract

**Objective:** There is increasing evidence for altered resting state functional connectivity (rsFC) in adolescents with disruptive behavior. Despite considerable ongoing behavioral research suggesting also important differences relating to reactive and proactive aggression, the corresponding rsFC correlates have not been studied to date. We therefore examined associations between these aggression subtypes along with subdimensions of callous-unemotional (CU) traits and rsFC using predefined seeds in aggression-related salience network (SN) and default mode network (DMN).

**Method:** Aggression subtype-specific whole-brain rsFC of SN and DMN seeds was investigated in a resting state sequence (mean acquisition time = 8 min 25 sec) acquired from 207 children and adolescents of both sexes aged 8 - 18 years (mean age (SD) = 13.30 (2.60) years; range = 8.02 – 18.35) in a multi-center study. One hundred eighteen individuals exhibited disruptive behavior (conduct disorder/oppositional defiant disorder) with different levels of comorbid ADHD symptoms, 89 were healthy.

**Results:** Compared to healthy controls, cases demonstrated reduced DMN and – after controlling for ADHD scores – SN seed-based rsFC with left hemispheric frontal clusters. We found increased and distinct aggression-subtype specific rsFC patterns. Specifically, reactive and proactive aggression correlated with distinct SN and DMN seed-based rsFC patterns. CU dimensions led to different DMN and SN rsFC with clusters including frontal, parietal, and cingulate areas.

**Conclusions:** This first study investigating reactive and proactive aggression along with CU dimensions reveals new subtype-specific whole-brain rsFC patterns in brain regions linked to processes like emotion, empathy, moral, and cognitive control.

## Introduction

Oppositional defiant disorder (ODD) and conduct disorder (CD) are among the most common psychiatric disorders in childhood and adolescence (Polanczyk, Salum, Sugaya, Caye, & Rohde, 2015), characterized by angry and vindictive behaviors, and violating rules, norms, and rights, respectively (American Psychiatric Association, 2013). Comorbid attention-deficit/hyperactivity disorder (ADHD) symptoms are frequently present (Waschbusch, 2002) with overlapping neural deficits in prefrontal and limbic areas (Puiu et al., 2018). Recent studies of resting state fMRI (rs-fMRI) in adolescents with CD have mainly reported reduced resting state functional connectivity (rsFC) or activity. Affected brain regions include the amygdala and insula as parts of the salience network (SN) (Aghajani et al., 2016, 2017; Zhou, Yao, Fairchild, Zhang, & Wang, 2015), and areas of the default mode network (DMN) (Broulidakis et al., 2016; Lu, Zhou, Wang, Xiang, & Yuan, 2017; Lu, Zhou, Zhang, Wang, & Yuan, 2017; Lu et al., 2015; Zhou et al., 2016). Connectivity between the core DMN regions anterior medial prefrontal cortex (amPFC) and the posterior cingulate cortex (PCC) was reduced in male adolescents with CD compared to healthy controls after controlling for ADHD symptoms, as ADHD symptoms correlated positively with DMN rsFC (Broulidakis et al., 2016). Other recent studies have also reported reduced DMN rsFC in male adolescents with CD compared to healthy controls (Zhou et al., 2016). Higher ADHD scores, however, related to increased rsFC density (Lu, Zhou, Wang, et al., 2017) and alterations (Lu, Zhou, Zhang, et al., 2017; Pu et al., 2017; Uytun et al., 2016) in DMN areas. Critically, these rs-fMRI studies were mostly limited to male adolescents with CD, and further distinct manifestations of aggression including reactive and proactive aggression (RA/PA) (Dodge & Coie, 1987) largely neglected.

Despite considerable research on a behavioral level on the importance of the differentiation between impulsive RA, and instrumental, PA behavior – with evidence for both being correlated (Polman, Orobio De Castro, Koops, Van Boxtel, & Merk, 2007) and also relating to different behavioral correlates (Fite, Stoppelbein, & Greening, 2009) – no rsFC analysis has addressed the role of RA and PA in children and adolescents with disruptive behavior to date.

CU traits are reported in 25-30% of children with an early onset of CD symptoms (Frick, 2016) and can be characterized by callousness, uncaring, and unemotional subdimensions (Essau, Sasagawa, & Frick, 2006; Pechorro, Ray, Gonçalves, & Jesus, 2017). To date, only few rsFC studies have evaluated CU subdimensions in children and adolescents. While the anticorrelation between the DMN and fronto-parietal network decreased with higher CU-related traits (Pu et al., 2017), anterior DMN rsFC increased with higher scores on the CU dimension (Cohn et al., 2015), and DMN rs-fMRI parameters increased with higher CU-related total scores and interpersonal/affective traits (Thijssen & Kiehl, 2017). In male youths with CD, interpersonal traits correlated with distinct amygdala subregional rsFC with clusters including SN and DMN regions (Aghajani et al., 2016). Compared to male youths with CD and lower CU traits or healthy controls, juveniles with CD and higher CU total scores showed increased amygdala subregional rsFC with a cluster including frontal and DMN regions (Aghajani et al., 2017). However, most rsFC studies to date restricted analyses to males and have not included healthy controls, children, and further aggression dimensions.

This is the first study of fMRI rsFC in boys and girls with disruptive behavior including diagnoses of CD and ODD that takes RA and PA along with CU dimensions into account. We applied a frequently used seed-based approach (Aghajani et al., 2016, 2017; Pujol et al., 2012; Uytun et al., 2016). Based on the findings of altered rsFC of the DMN (Broulidakis et al., 2016; Lu, Zhou, Wang, et al., 2017; Lu et al., 2015; Zhou et al., 2016) and SN (Aghajani et al., 2016, 2017), we defined regions of interest (ROIs) in these brain areas and expected to find reduced rsFC in aggressive cases compared to healthy controls. As RA and PA have not been addressed yet, we analyzed seed-based rsFC of amygdala and anterior insula (Blair, 2016; Fanning, Keedy, Berman, Lee, & Coccaro, 2017; Lozier, Cardinale, VanMeter, & Marsh, 2014). Further, we expected to find distinct patterns for different CU dimensions (Aghajani et al., 2016, 2017; Cohn et al., 2015) and increased rsFC with higher ADHD scores (Broulidakis et al., 2016; Lu, Zhou, Wang, et al., 2017).

## Method

### Participants

Participants in the current study were part of the joint EU-MATRICS and EU-Aggressotype project. Children and adolescents aged 8-18 years were recruited from resident hospitals, ambulatories, and eligible (boarding) schools. A total of 207 (*n* = 150 males) cases (*n* = 118) and healthy controls (*n* = 89) were included from nine different sites in Europe (mean age = 13.30 ± 2.60 years). As the main goal was to conduct aggression subtype-specific analyses, recruitment focused on including cases presenting with a diagnosis of CD and/or ODD and/or aggression scores in a clinical range (T > 70) according to the Child Behavior Checklist (CBCL), Youth Self Report (YSR), or Teacher Report Form (TRF) (Bordin et al., 2013). Controls were not allowed to have clinical aggression scores (T > 70) or a DSM-diagnosis. Further exclusion criteria for all participants were contraindications for MRI scanning (i.e., braces, metal parts) and insufficient intellectual and cognitive functioning. Participants and their parents or legal representatives gave written informed consent. Each site obtained ethical approval separately.

### Clinical Assessments

The semi-structured interview Kiddie-Schedule for Affective Disorders and Schizophrenia, present and lifetime version (K-SADS-PL) (Kaufman et al., 1997) was used to assess diagnostic criteria for all participants by trained psychologists or interns based on the reports of participants and their parents interviewed separately. The self-reported Reactive Proactive Aggression Questionnaire (Raine et al., 2006) measured RA and PA forms of aggression. To assess CU traits, parents filled out the Inventory of Callous-Unemotional traits (ICU) (Essau et al., 2006; Kimonis et al., 2008) consisting of three subscales assessing callousness, uncaring, and unemotional behaviors. ADHD symptoms were evaluated using the inattention, hyperactivity, and impulsivity counts of the K-SADS-PL (Kaufman et al., 1997). Further details are provided in the Supplemental Information.

### Image Acquisition and Preprocessing

For data acquisition, six sites used Siemens 3 Tesla (T) scanners, two sites Philips 3T scanners, and one site a GE 3T scanner (see Supplemental Table S1 and Table S2). T1-weighted anatomical scans with largely similar parameters across sites (see Supplemental Table S1) were used to include white matter and cerebrospinal fluid parameters as confound regressors during temporal preprocessing. T2*-weighted echo-planar resting state functional imaging was performed with predominantly similar parameters across sites (TR 2.45s or less, at least 32 slices; see Supplemental Table S2). During an average acquisition time of 8 min 25 sec, participants were instructed to lie still, look at a white crosshair presented against a black background, and let their mind wander by not thinking about anything in particular. Standard preprocessing steps were applied using SPM12 (Welcome Trust Centre for Neuroimaging, UCL, United Kingdom; http://www.fil.ion.ucl.ac.uk/spm) and the SPM-based CONN toolbox v17.b (http://www.nitrc.org/projects/conn). Since our cases presented with externalizing disorders including comorbid ADHD symptoms, we used a threshold for excessive motion recently applied in rs-fMRI analyses in adolescents with ADHD (von Rhein et al., 2016). This led to the exclusion of ten cases with a root mean square framewise displacement (RMS-FD) of > 0.95 mm. Sensitivity analyses applying an even more conservative threshold are provided in the Supplement. Twelve participants were excluded due to missing or insufficient quality of structural scans, 14 individuals based on image artifacts.

### Regions of Interest

Using MarsBar toolbox (v0.44) (http://marsbar.sourceforge.net), PCC and amPFC were centered on coordinates recently used (Broulidakis et al., 2016) provided by Andrews-Hanna et al. (Andrews-Hanna, Reidler, Sepulcre, Poulin, & Buckner, 2010). Additionally, bilateral amygdala and bilateral anterior insula as part of the SN were derived from the Broadman-atlas (http://fmri.wfubmc.edu/software/pickatlas).

### Functional Connectivity Analysis

Seed-based rsFC analyses were performed using CONN toolbox. First-level analysis computed Pearson’s correlation coefficients between the time course of previously denoised BOLD-signals from a seed and whole-brain voxel clusters. After Fisher’s transformation to normally distributed z-scores, general linear model (GLM) analyses were computed. Second-level analysis included random-effects analysis of covariance for group comparisons, with further analyses adding ADHD symptoms as additional covariates of no interest based on previous reports on the importance of considering ADHD symptoms to differentiate rsFC of cases and controls (Broulidakis et al., 2016; Uytun et al., 2016). Linear regressions separately tested the association between RA and PA and CU total score and dimensions on rsFC within cases. Besides site added as dummy-coded covariate of no interest in second-level analyses, we additionally controlled for age, sex, IQ, medication, and handedness given previous reports on possible influences for instance on rsFC of the DMN (Mak et al., 2017). Results of the seed-based analyses are reported at a statistical threshold of *p* < .001, *p*-FWE < .008 cluster-level corrected (=0.05/6, using additional Bonferroni corrections for number of seeds).

## Results

### Sample Characteristics

Out of 118 cases, 48 had a diagnosis of ODD, 25 of CD plus ODD, and seven of CD. Seventy-seven cases presented with a clinically relevant score (T > 70) on aggression or rule-breaking behavior subscales of the CBCL, and 41 cases on both subscales. Thirty-eight cases had an aggression score in the clinical range but no DSM-diagnosis (Table 1). While cases and controls were matched regarding age and handedness, cases consisted of more males than females, exhibited a lower IQ than healthy controls, and showed a wide distribution of RA and PA levels, CU traits, and ADHD symptoms. The proportion of controls relative to cases was not consistent across participating sites. For the distribution of diagnoses, aggression scores, medication, and demographic variables across sites, see Supplemental Table S3.

**Table 1.** Sample Characteristics

| Characteristic | Cases<br>(*n* = 118) | HC<br>(*n* = 89) | Test statistic | *p* |
| --- | --- | --- | --- | --- |
| Age (Years) | 13.23 ± 2.68 | 13.40 ± 2.49 | *t*(205) = -0.45 | 0.65 |
| Sex, m/f | 99/19 | 51/38 | $\chi^2 = 17.98$ | < 0.001 |
| IQ<sup>a</sup> | 100.78 ± 11.00 | 106.64 ± 10.42 | *t*(195) = -3.81 | < 0.001 |
| Handedness, left/right | 16/95 | 10/77 | $\chi^2 = 0.37$ | 0.55 |
| CD plus ODD Diagnosis<sup>b</sup> | 25 |  |  |  |
| ODD Diagnosis<sup>b</sup> | 48 |  |  |  |
| CD Diagnosis<sup>b</sup> | 7 |  |  |  |
| ADHD Diagnosis<sup>b</sup> | 29 |  |  |  |
| CBCL – Aggression T-score | 74.46 ± 10.10 |  |  |  |
| CBCL – Rule-breaking T-score | 69.00 ± 12.14 |  |  |  |
| K-SADS - Inattention | 3.33 ± 2.91 |  |  |  |
| K-SADS - Hyperactivity | 1.66 ± 1.91 |  |  |  |
| K-SADS - Impulsivity | 1.08 ± 1.20 |  |  |  |
| ICU – Total Score | 33.68 ± 10.16 | 21.00 ± 8.70 | *t*(196) = -9.44 | < 0.001 |
| ICU – Callousness | 12.00 ± 6.11 | 4.00 ± 3.44 | *U* = 2278.00 | < 0.001 |
| ICU – Uncaring | 17.00 ± 3.93 | 10.41 ± 5.07 | *U* = 2445.00 | < 0.001 |
| ICU – Unemotional | 7.17 ± 3.31 | 5.22 ± 2.75 | *t*(189) = -4.37 | < 0.001 |
| RPQ – Reactive Aggression | 12.55 ± 5.09 | 5.00 ± 3.48 | *U* = 1296.50 | < 0.001 |
| RPQ – Proactive Aggression | 3.00 ± 5.01 | 0.82 ± 1.45 | $U = 1807.50$ | < 0.001 |
| Medication use<sup>c</sup> | 70 |  |  |  |
| - Stimulants | 52 |  |  |  |
| - Neuroleptics | 18 |  |  |  |
| - Antidepressants | 2 |  |  |  |
| Mean RMS-FD | 0.12 ± 0.17 | 0.09 ± 0.18 | $U = 4134.50$ | <0.01 |

Values are means or in case of non-normal distribution medians ± SD, or counts.
HC, healthy controls; ADHD, attention-deficit hyperactivity disorder; CD, conduct disorder; ODD, oppositional defiant disorder; K-SADS, Kiddie Schedule for Affective Disorders and Schizophrenia; ICU, Inventory of Callous-Unemotional traits, parent report; RPQ, Reactive and Proactive aggression Questionnaire; CBCL; Child Behavior Checklist; RMS-FD, root mean square framewise displacement.
<sup>a</sup>IQ estimated according to four sub-tests derived from the Wechsler Intelligence Scale for Children IV.
<sup>b</sup>Diagnoses derived from the Kiddie-Schedule for Affective Disorders and Schizophrenia, present and lifetime version.
<sup>c</sup>Medication use according to parental or clinical report.

### Group Differences in Functional Connectivity

We did not find group differences in our primary analyses, which included the control for age, sex, IQ, medication, and handedness besides site. When we controlled for site only, cases demonstrated a reduced DMN (PCC seed) connectivity with a cluster including left frontal pole (*t*(197) = 5.46, cluster-size *p*-FWE < .008, peak *p*-uncorrected < .00001, *β* =.10) (Figure 1, Supplemental Table S4), surviving the control for site, age, sex, and IQ at a less conservative significance threshold (cluster-size *p*-FWE < .05) and a more stringent excessive motion criterion (sensitivity analyses, Supplemental Table S9). After taking ADHD symptoms into account, cases showed a diminished left hemispheric rsFC of SN (anterior insula seed) with a cluster extending from orbitofrontal cortex (OFC) to frontal pole compared to controls (*t*(194) = 5.07, cluster-size *p*-FWE < .008, peak *p*-uncorrected < .00001, *β* = .10) (Figure 1, Supplemental Table S4). This group difference survived correction for site, age, sex, IQ, and handedness. Subsequent analysis within cases revealed a positive correlation of ADHD inattention and hyperactivity counts and SN (left anterior insula seed) rsFC, however at a lower significance threshold (*t*(85) > 5.27, all cluster-size *p*-FDR < .05, peak *p*-uncorrected < .00001, *β* = .04-.09) (Figure 1). In order to explore the influence of the additional covariates sex, IQ, medication, and handedness on group differences in rsFC, we computed sensitivity analyses, which showed that none of these covariates had a significant influence on rsFC patterns (all *p* > .05).

**Figure 1.**
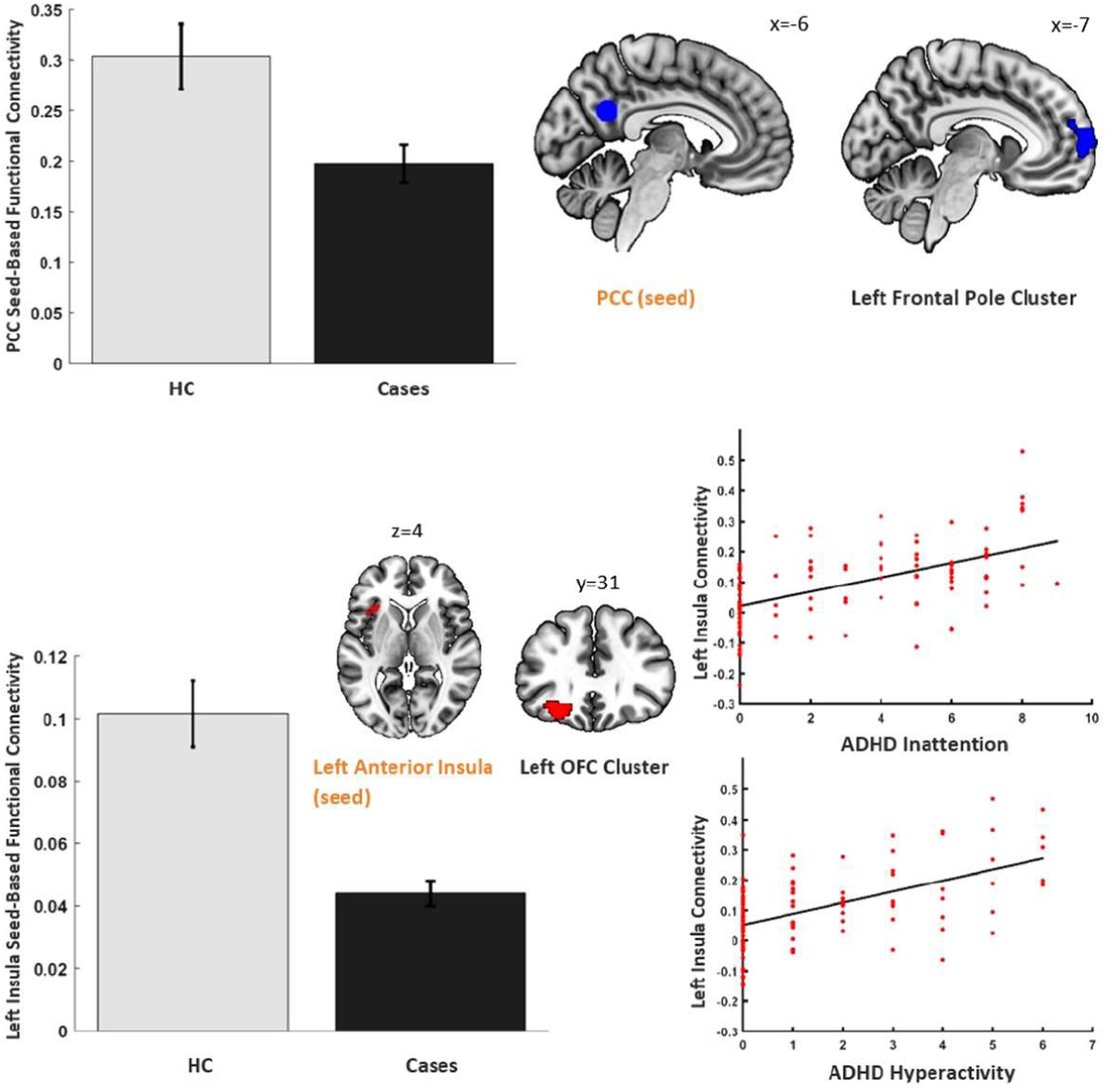
The bar charts show reduced seed-based functional connectivity patterns for cases compared to healthy control (HC) subjects. The scatterplots depict the main effect of ADHD inattention and hyperactivity counts within cases on SN seed-based connectivity with a left frontal cluster.

### Functional Connectivity Correlates of Reactive versus Proactive Aggression

RA and PA yielded distinct and increasing rsFC patterns of DMN and SN seeds within cases (Figure 2, Supplemental Table S6). PA scores related to SN (left amygdala seed) connectivity with a precuneal cluster. In contrast, cases with higher RA scores related to increased DMN (PCC seed) connectivity with a cluster extending from left parahippocampal gyrus to left inferior temporal gyrus. Furthermore, RA scores correlated with SN (left amygdala seed) rsFC with a cluster extending from left lateral occipital cortex to the precuneus. RA scores also related to SN (right anterior insula seed) rsFC with a cluster in the right caudate nucleus (all cluster-size *p*-FWE < .008, *β* = .04-.05). The findings remained significant at an uncorrected p<.01 threshold after the more stringent excessive motion criterion (sensitivity analyses, Supplemental Table S10).

**Figure 2.**
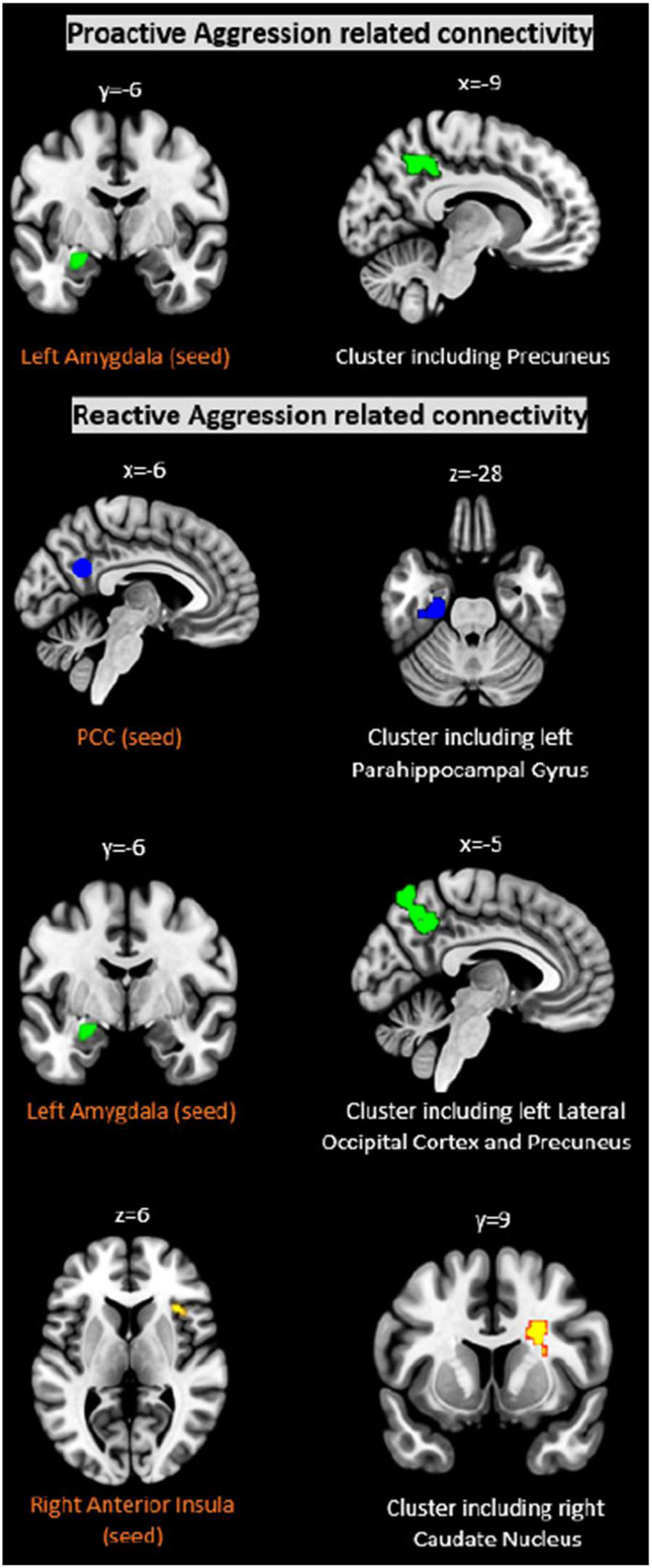
Distinct RA and PA-related connectivity patterns within cases (FWE-corrected, p < 0.008). We only observed one partly overlapping left amygdala seed-based pattern related to both RA and PA scores, yet with differing correlated clusters.

### Functional Connectivity Correlates of Specific CU Dimensions

The different CU dimensions were positively associated with distinct seed-based rsFC patterns within cases (Figure 3, Supplemental Table S7). Callousness-related DMN seed-based connectivity patterns included clusters in the right precentral gyrus. Clusters correlating with amPFC seed extended to the parietal and occipital areas and the cluster for PCC seed further included cingulate regions. Uncaring and unemotional behavior scores related to SN (left anterior insula seed) rsFC with precuneal and cingulate clusters. Uncaring-related clusters extended to right central gyrus, while unemotional-related clusters expanded to left occipital and parietal areas, such as the angular gyrus. Uncaring behavior scores also related to SN (right anterior insula seed) rsFC with a cluster in the left central gyrus and to DMN (amPFC seed) rsFC with right hemispheric cerebellar regions (all cluster-size *p*-FWE < .008, *β* = −.06−.08) (Figure 3, Supplemental Table S7).

**Figure 3.**
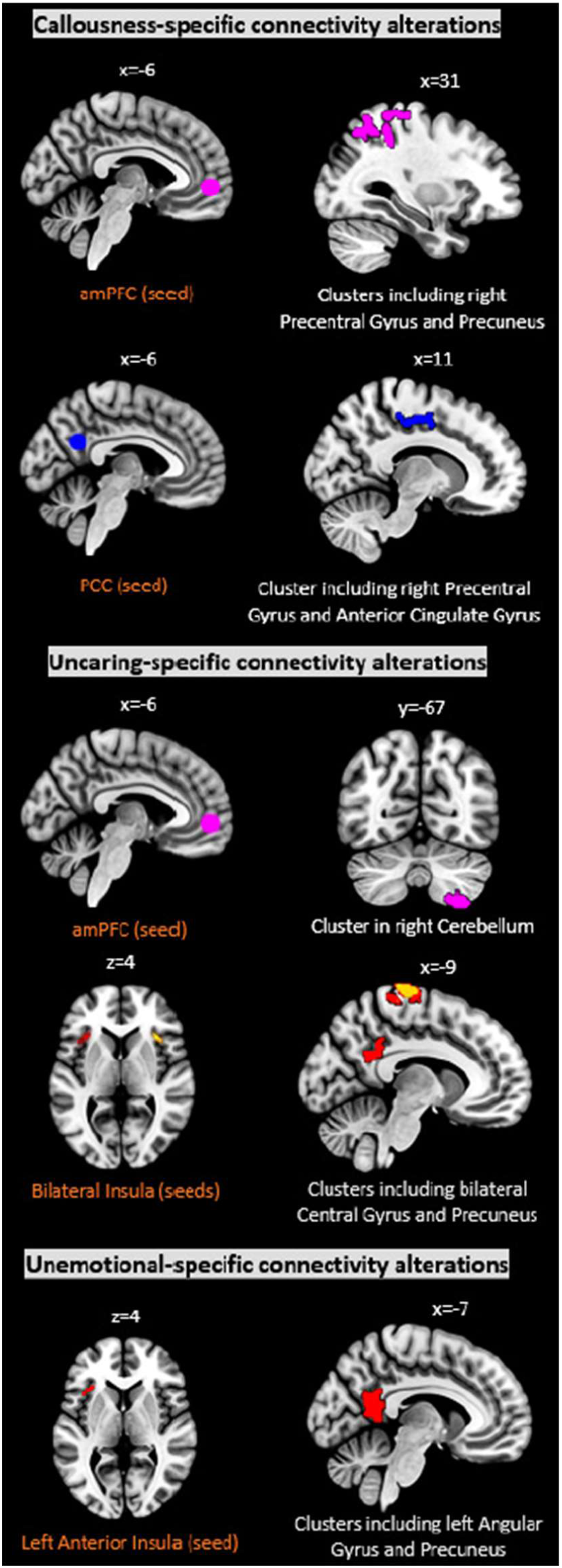
Differing rsFC patterns related to scores on the CU subdimensions callousness, uncaring, and unemotional within cases (FWE-corrected, p < 0.008).

## Discussion

The present multi-center study investigated aggression subtype-specific rsFC in a sizeable sample of children and adolescents with disruptive behavior including diagnoses of CD and ODD. Cases exhibited diminished DMN (PCC seed) rsFC with a frontal cluster including the left frontal pole compared to healthy controls. The additional control for ADHD symptoms further identified decreased SN (left anterior insula seed) rsFC with a left hemispheric frontal cluster. We found distinct and increasing aggression subtype-specific rsFC patterns within cases in brain regions linked to processes like emotion and empathy. PA scores related to SN connectivity with a cluster including the precuneus, and RA scores related to distinct DMN and SN seed-based rsFC patterns with (para-)limbic and precuneal clusters. Callousness and uncaring behavior scores related to different DMN and SN rsFC with voxel clusters including frontal, parietal, cingulate, precuneal, and cerebellar areas. Both uncaring and unemotional behavior scores correlated with SN rsFC with precuneal and cingulate clusters, which extended to distinct brain areas including the central and angular gyrus, respectively.

Abnormal rsFC patterns for the PCC have been demonstrated in male adolescents with CD (Broulidakis et al., 2016; Lu, Zhou, Wang, et al., 2017; Zhou et al., 2016) and may suggest impaired self-referential processes (Andrews-Hanna et al., 2010). As observed group differences only survived post hoc analysis when other covariates were not controlled for or a lower threshold was applied, cautious interpretation is advised. Our results support the crucial role of comorbid ADHD symptoms previously shown in male adolescents with CD within the DMN (Broulidakis et al., 2016), and extend analysis to DMN and SN seed-based rsFC with whole-brain voxel clusters. Diminished SN rsFC with left hemispheric clusters extending from the OFC to the frontal pole in cases compared to controls were only demonstrated when considering ADHD scores. In line with previous rs-fMRI studies (Broulidakis et al., 2016; Lu, Zhou, Wang, et al., 2017; Pu et al., 2017), cases exhibited an ADHD scores-related increase in rsFC and further corroborates the recent finding of overlapping deficit functioning in ADHD and disruptive behavior disorders (Puiu et al., 2018).

In line with our hypothesis, RA and PA related to distinct SN and DMN seed-based rsFC patterns in brain areas implicated in emotion, empathy, and cognitive control. RA scores correlated with increased rsFC within paralimbic and limbic regions shown to function abnormally in psychopathy (Espinoza et al., 2018), which partly corroborates the previously reported behavioral overlap of RA and PA with CU-related traits (Kimonis et al., 2008; Pechorro et al., 2017). The cluster involved in SN (right anterior insula seed) rsFC included the right caudate nucleus, a region implicated in integration of performance and cognitive control (Brovelli, Nazarian, Meunier, & Boussaoud, 2011). Neural circuits including the anterior insula have been linked to responses to frustration and perceived social provocations (Blair, 2016). Moreover, both aggression subtypes related to increased SN (left amygdala seed) rsFC with a cluster including the precuneus. Impaired amygdala functioning seems involved in both a neural threat circuitry related to a higher risk for reactive aggression and in moral behavior increasing the risk for PA (Blair, 2010). Further, rsFC of the precuneus was found to be related to impulsivity in a recent study (Lu, Zhou, Wang, et al., 2017), which is relevant given the association of impulsivity and RA (Fite et al., 2009). However, the cluster implicated in SN rsFC extended to distinct paralimbic and occipital areas for RA and PA, which underpins their differentiation in line with behavioral findings (Fite et al., 2009).

The reported different rsFC patterns for the distinct CU dimensions corroborate previous neural (Aghajani et al., 2016, 2017; Cohn et al., 2015; Espinoza et al., 2018; Philippi et al., 2015) and behavioral (Pechorro et al., 2017) findings. Affected brain areas extended from (para-)limbic regions implicated in adult psychopathy (Espinoza et al., 2018) to frontal, parietal, and cingulate areas. These regions have been linked to emotion, empathy, moral, and self-referential processes which are impaired in youth with disruptive behaviors (Blair, Veroude, & Buitelaar, 2016). In line with rsFC studies in adolescents (Aghajani et al., 2016, 2017; Cohn et al., 2015) and adults (Philippi et al., 2015; Tang et al., 2016), cases exhibited increases in rsFC with higher CU traits. Callousness and uncaring behavior were related to rsFC patterns including clusters in the precentral gyrus, which has been previously linked to altered rsFC in adolescents with CU-related traits (Cohn et al., 2015; Pu et al., 2017) and psychopathic adults (Espinoza et al., 2018; Korponay et al., 2017; Philippi et al., 2015; Tang et al., 2016). A recent meta-analytic review found that the right anterior insula was implicated in the evaluation of feelings and the left anterior insula in the expression of anger (Lindquist, Wager, Kober, Bliss-Moreau, & Barrett, 2012). The DMN seed-based rsFC patterns reported here might suggest uncaring and callousness dimension-specific alterations in (affective) self-referential processes (Andrews-Hanna et al., 2010). Altered rsFC in the DMN was reported in children and adolescents with CU-related traits (Cohn et al., 2015; Pu et al., 2017) and in adult psychopathy (Pujol et al., 2012; Tang, Jiang, Liao, Wang, & Luo, 2013; Tang et al., 2016). The overlapping SN rsFC with a cluster in precuneal and cingulate areas for uncaring and unemotional behavior scores may reflect their behavioral correlation (Kimonis et al., 2008). The precuneus as part of the DMN contributed to classifying adults with antisocial personality disorder (Tang et al., 2013). Altered insular and cingulate functioning has been related to moral reasoning in adult psychopathy (Griffiths & Jalava, 2017). We did not find overlapping rsFC between CU dimensions and RA or PA despite well-known behavioral commonalities (Kimonis et al., 2008; Pechorro et al., 2017) and the behavioral correlation of PA scores with CU traits in our sample. Surprisingly, no rsFC pattern included the amygdala contrary to previous reports (Aghajani et al., 2016, 2017; Salekin, 2017), which may provide support for recent findings that suggest neural alterations beyond (para-)limbic regions in adult psychopathy (Griffiths & Jalava, 2017).

There are some limitations in the present study. Firstly, 38 aggressive cases without a DSM-diagnosis of CD and/or ODD exhibited lower PA scores compared to cases with a DSM-diagnosis. However, subsequent sensitivity analyses led to comparable results of case-control group comparisons in rsFC when excluding these cases without a diagnosis. Secondly, we did not consider self-reports of CU traits. Yet, a recent study reported a higher criterion validity for parent reported ICU compared to self- and teacher reports (Docherty, Boxer, Huesmann, O’Brien, & Bushman, 2017). Thirdly, the distribution of cases and controls across sites was not balanced and case and control groups not matched regarding sex, IQ, and number of participants. Thus, we conducted sensitivity analyses, which showed no significant influence of sex, IQ, medication, and handedness on rsFC patterns of cases compared to controls. Fourthly, data for the current study was collected at different sites with varying scanner manufacturers and partly deviating scan parameters, which affected data homogeneity and limited our study power. However, the larger sample size enabled by the multi-center design might have increased the reliability and generalizability of our results.

Taken together, in the present study children and adolescents with disruptive behavior exhibited decreased rsFC with left hemispheric frontal voxel clusters. More importantly, cases demonstrated an aggression subtype-specific increase in DMN and SN seed-based whole-brain rsFC with clusters beyond (para-)limbic areas in frontal, parietal, and cingulate regions related to emotion and empathy. By evaluating the effect of RA and PA along with CU dimensions, we have extended previous research in mainly male adolescents with CD and limited exploration of distinct manifestations of aggression. Our aggression subtype-specific findings provide support for a more precise diagnostic specification of aggression-related disorders. Particularly treatment of children and adolescents with disruptive behavior may be improved through careful exploration of distinct aggression subtypes and a better understanding of neural correlates. Further, our results may indicate developmental trajectories with some of the observed brain areas relating to adult psychopathy.

## Supporting information

Supplement_2ndManuscript_Werhahn

## Acknowledgments

This project has received funding from the European Union’s Seventh Framework Programme for research, technological development and demonstration under grant agreement 602805 (Aggressotype) and no 603016 (MATRICS). This manuscript reflects only the author’s view and the European Union is not liable for any use that may be made of the information contained herein. MC is currently funded by the Medical Research Council UK (grant MR/M013588). TB served in an advisory or consultancy role for Actelion, Hexal Pharma, Lilly, Lundbeck, Medice, Neurim Pharmaceuticals, Novartis, Shire. He received conference support or speaker’s fee by Lilly, Medice, Novartis and Shire. He has been involved in clinical trials conducted by Shire & Viforpharma. He received royalties from Hogrefe, Kohlhammer, CIP Medien, Oxford University Press. CA has been a consultant to or has received honoraria or grants from Acadia, Ambrosseti, Gedeon Richter, Janssen Cilag, Lundbeck, Merck, Otsuka, Roche, Servier, Shire, Schering Plough, Sumitomo Dainippon Pharma, Sunovion and Takeda. DB serves as an unpaid scientific advisor for an EU-funded neurofeedback trial unrelated to the present work. BF receives funding from a personal Vici grant of the Dutch Organisation for Scientific Research (grant 016 130 669) and from the Dutch National Science Agenda for the NWANeurolabNL project (grant 400 17 602). She received educational speaking fees from Shire and Medicine. SW has received lecture honoria from Lilly, Opopharma/Shire in the last five years. She received royalties from Hogrefe, Kohlhammer, Springer, Beltz. Outside professional activities and interests are declared under the link of the University of Zurich www.uzh.ch/prof/ssl-dir/interessenbindungen/client/web/. The present work is unrelated to the above grants and relationships. The authors do not have potential conflicts of interest. The authors express their gratitude to all participating families and thankfully acknowledge the contribution of Yu Jin Ressel, Noemi Signer, Aline Pfister, Georg Grön, Kathrin Brändle.

## Key points

- Increasing evidence suggests mainly reduced resting state functional connectivity (rsFC) in adolescents with disruptive behavior in regions of the default mode network (DMN) and salience network (SN).
- While some rsFC studies evaluated the neural correlates of CU traits, reactive and proactive (RA/PA) forms of aggression have been neglected to date.
- We investigated DMN and SN seed-based rsFC in a large multicenter sample of children and adolescents, considering RA/PA behaviors along with CU traits.
- We found reduced rsFC in aggressive cases compared to controls in frontal clusters, with one pattern depending on the additional control for ADHD scores.
- Within cases, we found subtype-specific whole-brain rsFC patterns in brain regions previously linked to emotion, empathy, moral, and cognitive control.

