## Supplement_2ndManuscript_Werhahn for "Aggression Subtypes Relate to Distinct Resting State Functional Connectivity in Disruptive Children and Adolescents"

***Supplemental Information***

**Supplementary Methods and Materials**

**Participants**

The current analyses included 207 children and adolescents. One hundred eighteen cases exhibited disruptive behavior (mean age = 13.23, SD = 2.68) and 86 were age- and handedness-matched healthy controls (mean age = 13.40, SD = 2.49). Within cases, 25 had a DSM-5 diagnosis of conduct disorder (CD) and of oppositional defiant disorder (ODD), 48 of ODD, seven of CD, 77 cases (additionally) presented with an aggression score in the clinical range (T > 70) on the aggression or rule-breaking behavior subscales according to the Child Behavior Checklist (CBCL), and 41 cases on both subscales. Thirty-eight cases had an aggression score in the clinical range but no DSM-diagnosis. Out of 29 cases with comorbid ADHD diagnosis, 15 cases had a diagnosis of ODD only and one case of CD only, and 8 had a diagnosis of ODD plus CD. None of our controls met a diagnosis of ADHD nor ADHD inattention or hyperactivity/impulsivity syndromes according to the K-SADS. Participants were recruited at nine different sites in Europe (Radboud University Medical Center and the Donders Center for Cognitive Neuroimaging, Nijmegen, The Netherlands; Department of Neuroscience, University Medical Centre Groningen, The Netherlands; Central Institute of Mental Health, Medical Faculty, Mannheim/Heidelberg University, Mannheim, Germany; University of Ulm, Department of Child and Adolescent Psychiatry/Psychotherapy and Department of Psychiatry III, Ulm, Germany; Department of Child Psychiatry, and the Centre for Neuroimaging Sciences, Institute of Psychiatry, Psychology and Neuroscience, King`s College London, London, England; Institut d'Investigacions Biomèdiques August Pi i Sunyer and Hospital Clinic de Barcelona, Barcelona, Spain; Instituto de Investigación Sanitaria Gregorio Marañón, Child and Adolescent Psychiatry Department and Ruber International Hospital Madrid, Madrid, Spain; Department of Child and Adolescent Psychiatry and Psychotherapy, University Zurich and MR center, Psychiatric University Hospital, Zurich, Switzerland; IRCCS Santa Lucia Foundation, Rome, Italy). Recruitment took place at resident hospitals, ambulatories, and eligible (boarding) schools.

Besides a DSM-5 diagnosis of CD, ODD, and/or an aggression or rule-breaking behavior subscale score in a clinical range (T > 70) according to the CBCL, Youth Self Report (YSR), or Teacher Report Form (TRF), further inclusion criteria for the case group were no medication or a stable medication at least for two months. Exclusion criterion was a primary DSM-5 diagnosis of depression, anxiety, psychosis, or bipolar disorder for cases, and for healthy controls a DSM-5 diagnosis or clinically relevant scores in the CBCL, YSR, or TRF. Further exclusion criteria for all participants were an IQ score < 80 as measured by the Wechsler Intelligence Scale for Children, Fourth Edition (WISC-IV), and common contraindications for MRI scanning, such as braces or metal parts. Additionally, an anxiety score >8 on a Visual Analogue Scale ranging from 1 to 10 before scanning also lead to exclusion, as anxiety caused by the scanner could impair data quality and alter neural activity (1). Participants had sufficient native language skills according to the assessing country. All sites obtained ethical approval separately. After participants received information about the study procedure, participants and their parents or legal representatives gave written informed consent.

Before conducting further analyses, we excluded 12 participants (11 cases, 1 control) due to missing (*n* = 2) or insufficient quality of the T1 weighted anatomical scans (*n* = 10) from our original data set of resting state sequences from 141 cases and 92 healthy controls. Moreover, 12 cases and 2 controls were excluded due to image artifacts and 10 cases due to excessive motion. Finally, 118 cases and 89 controls, both males (*n* = 150) and females (*n* = 57) aged 8 to 18 years (M = 13.30, SD = 2.60) were included in the resting state functional connectivity (rsFC) analysis.

**Assessment Tools and Study Procedure**

Diagnostic assessments and MR measurements took place on different dates to minimize burden for the participants. During the first appointment, parents/primary caregivers and children/youths were assessed by trained (clinical) psychologists or interns separately with the semi-structured interview Kiddie-Schedule for Affective Disorders and Schizophrenia, present and lifetime version K-SADS (2). The full supplementary module of the specific disorder followed positively answered questions. Diagnoses resulted from self- and parent-reports. If not already completed and brought to the assessment, parents/primary caregivers and children/youths answered a questionnaire package. The Child Behavior Checklist (CBCL) is a parent-report questionnaire on which the child or adolescent is rated on various behavioral and emotional problems (3). The Swanson, Nolan, and Pelham teacher and parent rating scale (SNAP-IV) contains 26 items to measure attention deficit disorder (ADHD) and oppositional defiant disorder (ODD) symptoms from childhood to young adulthood (4). The Inventory of Callous-Unemotional traits (ICU) is a 24-item questionnaire ranging from 0 (not at all true) to 3 (definitely true), and forming three subscales to assess callous and unemotional traits: uncaring, callousness, and unemotional (5). The Reactive-Proactive Aggression Questionnaire (RPQ) is a 23-item self-report measure of the frequency (never: 0, sometimes: 1, often: 2) of reactive and proactive aggression (6). IQ was estimated according to four sub-tests derived from the Wechsler Intelligence Scale for Children (WISC-IV) (7): block design, similarities, vocabulary, and picture completion. Additionally, digit span sub-test was assessed. Further parts of test battery in the framework of the Aggressotype/MATRICS study were also conducted and included additional questionnaires (Modified Aggression Scale (8), Longterm difficulties self- and parent-report (9), Pubertal Developmental Scale (10), Antisocial Behavior Scale (11), Strengths and Difficulties Questionnaire (12), Youth Self Report (YSR) and Teacher Report Form (TRF) as further parts of Achenbach System of Empirically Based Assessment (3) as well as ICU (5) self- and teacher-report, cognitive tests (Probabilistic Reversal Learning Task (13) and three tasks from the Cambridge Neuropsychological Test Automated Battery (14): Emotion Recognition Task, Delayed Matching to Sample and Rapid Visual Information Sampling), and blood- or saliva-samples for biological analysis. After a short psychophysiological measurement (including measures of heart rate (pulse), skin conductance level, and EEG before, during, and after and emotional paradigm), children/youths were prepared for MR scanning in attendance of their parents/primary caregivers. They were presented with MR sounds, and visited a dummy scanner and/or watched a MR information video. If the participants or parent/primary caregiver reported an anxiety rating of >8 on a Visual Analogue Scale (VAS) ranging from 1 (no anxiety) to 10 (very high anxiety) related to entering the MR scanner, the participant was then excluded from further investigation.

If all inclusion criteria were fulfilled, MR scanning took place on another appointment. Participants answered a MR safety form and a short pre-scanning questionnaire (e.g., regarding tobacco and cannabis consummation, genetic diseases in the family) and then practiced the following fMRI tasks on a laptop: passive avoidance task (15), emotional matching task (16), and stop signal task (17). Participants gave a saliva sample right before MRI scanning. They were asked to use the toilet, and female pregnancy was ruled out by a urine pregnancy test. The first part of MRI scanning session included a T1-weighted anatomical image, the three fMRI tasks mentioned above, and the fMRI resting state sequence. For the resting state scan, participants were instructed to look at a white crosshair against a black background and let their mind wander. After a break that included the collection of another saliva sample and the assessment of a short questionnaire regarding the performance in the fMRI tasks, a T1-weighted anatomical scan was followed by two sequences of magnet resonance spectroscopy and diffusion tensor imaging. After scanning, the granted monetary reward and travel costs were reimbursed. Additionally, participants received a picture of their anatomical MR scan. The current study reports resting state fMRI and psychometric data.

**Data Acquisition and Preprocessing**

For image acquisition, six sites used a Siemens 3T scanner, two sites a Philips 3T scanner, and one site a GE 3T scanner. Table S1 and Table S2 show the site-specific scanner information.

| **TABLE S1.** Structural MRI scan parameters across sites. | | | | | | | |
| --- | --- | --- | --- | --- | --- | --- | --- |
| **Scanner** | **Site** | **TR/TE/T1 (ms)** | **Flip angle** | **Field of view** | **Matrix RL/AP/slices** | **Voxel size (mm)** | **Acceleration factor** |
| Siemens | Nijmegen | 2300/2.98/900 | 9 | 256 | 212/256/176 | 1.0x1.0x1.2 | 2 |
|  | Mannheim | 2300/2.96/900 | 9 | 256 | 212/256/176 | 1.0x1.0x1.2 | 2 |
|  | Ulm | 2300/2.96/900 | 9 | 256 | 212/256/176 | 1.0x1.0x1.2 | 2 |
|  | Barcelona | 2300/2.98/900 | 9 | 256 | 212/256/176 | 1.0x1.0x1.2 | 2 |
|  | Madrid | 2300/2.98/900 | 9 | 256 | 212/256/176 | 1.0x1.0x1.2 | 2 |
|  | Rome | 2080/2.86/900 | 9 | 256 | 212/256/176 | 1.0x1.0x1.2 | 2 |
| Philips | Groningen | 2450/3.11/900 | 8 | 270 | 256/232/170 | 1.0x1.0x1.0 | 1.8 |
|  | Zurich | 2300/3.11/900 | 9 | 270 | 256/232/170 | 1.0x1.0x1.0 | 1.8 |
| GE | London | 2300/3.02/400 | 11 | 270 | 256/256/196 | 1.0x1.0x1.2 | 1.75 |

| **TABLE S2.** Resting state functional MRI scan parameters across sites. | | | | | | |
| --- | --- | --- | --- | --- | --- | --- |
| **Scanner** | **Site** | **TR/TE1/TE2/TE3 (ms)** | **Number of slices** | **Slice scan order** | **Voxel size (mm)** | **Duration (min)** |
| Siemens | Nijmegen | 2300/12/28.4/44.8 | 33 | descending | 3.8x3.8x3.8 | 8:24 |
|  | Mannheim | 2300/12/29/46 | 33 | descending | 3.8x3.8x3.8 | 8:24 |
|  | Ulm | 2300/31 | 33 | descending | 3.8x3.8x3.8 | 8:23 |
|  | Barcelona | 2300/12 | 33 | descending | 3.8x3.8x3.8 | 8:21 |
|  | Madrid | 2300/13 | 36 | descending | 3.8x3.8x3.8 | 8:24 |
|  | Rome | 2080/30 | 32 | ascending | 3.0x3.0x2.5 | 7:38 |
| Philips | Groningen | 2450/8.01/22.02/36.02 | 45 | descending | 3.5x3.5x3.5 | 10:08 |
|  | Zurich | 2300/13/31/49 | 33 | descending | 3.75x3.75x3.79 | 7:51 |
| GE | London | 2300/11.8/31/48 | 33 | descending, interleaved | 3.45x3.45x4.20 | 8:15 |

Preprocessing steps in SPM12 (Welcome Trust Centre for Neuroimaging, UCL, UK; <http://www.fil.ion.ucl.ac.uk/spm>) included realignment and unwarping (Andersson et al., 2001), followed by slice timing. Subsequently, multi-echo resting state data was linearly weighted by its echo time (TE) using Matlab (The MathWorks, MA, USA): Y4 = (Y1 * TE1 / TEsum) + (Y2 * TE2 / TEsum) + (Y3 * TE3 / TEsum), where Y is the echo file and TE the echo time. Using the SPM-based CONN toolbox v17.b (18), functional normalization to the Montreal Neurological Institute (MNI) template brain was used to reduce variability across subjects (19), followed by identification of outliers with the implemented ART-toolbox used for scrubbing during denoising. For functional smoothing, we used a 6-mm full-width at half-maximum Gaussian kernel, and segmented and normalized structural images for inclusion of white matter (WM) and cerebrospinal fluid (CSF) parameters as confound regressors. The aCompCor strategy (20) implemented in CONN during denoising enabled reducing the effect of physiological and motion-related noise (18). After estimating principal components of the subject-specific WM and CSF confounds using a principal component analysis of their BOLD signal, aCompCor removed the estimated timeseries by adding them as regressors. Additionally, the movement parameters derived from realignment were added as regressors. After denoising, initial hemodynamic response function signal aberrations were removed. Compared to the global signal regression method, aCompCor shows a higher sensitivity and specificity regarding positive correlations, and comparable specific and sensitive anticorrelations (21). We applied temporal band-pass filtering (0.008 – 0.9 Hz) in order to circumvent the influence of low-frequency drifts and high-frequency noise including heart rate and respiration (18).

**Motion Censoring**

Since head motion can lead to changes in the BOLD signal (22) and our cases presented with externalizing disorders including comorbid ADHD symptoms, we used a threshold for excessive motion recently applied in fMRI resting state analyses in adolescents with ADHD (23). Thus, scans exceeding the threshold of >5% of the highest mean Root Mean Square (RMS) framewise displacement (FD) values (>0.95 mm) (24) were excluded from further analysis. Moreover, we decided to apply also an even more conservative threshold for the detection of functional outlier scans using Artifact Detection Tools (ART, www.nitrc.org/projects/artifact_detect) implemented in CONN (>3mm standard deviations from the observed global BOLD signal and >0.5mm composite scan-to-scan motion). During denoising, the aCompCor strategy (20) implemented in CONN removed these ART-detected movement parameters by adding them as regressors.

**Supplementary Results**

Table S3 shows the distribution of demographic characteristics, diagnoses, and clinically relevant aggression scores on behavioral measures across sites for our final sample including 118 cases and 89 healthy controls. For seed-based group comparisons and aggression-subtype specific analysis within cases, a statistical threshold of *p* < .001, *p* < .008 FWE cluster-level correction (= .05 / 6 using additional Bonferroni correction for number of seeds) for multiple comparisons was applied (see Table S4, Table S6, Table S7). We also conducted sensitivity analyses after exclusion of one site due to a small sample size for case group (*n* < 5) (Table S5). Moreover, after excluding further 10 cases and 5 controls exceeding the threshold of RMS-FD >0.5 mm, we run sensitivity analyses by rerunning the main seed-based analyses in cases (*N* = 108) compared to controls (*N* = 84) (see Tables S9) and within cases after correcting for all covariates of no interest (see Table S10 and Table S11). This resulted in comparable case-control differences in rsFC and new additional patterns within cases, while some aggression-related patterns within cases only survived at a slightly lower uncorrected height threshold of *p* < .01 (see Table S10 and Table S11).

The average RMS-FD for cases was 0.12mm (*SD* = 0.17mm) and for controls 0.09 (*SD* = 0.18mm). In the whole sample, mean RMS-FD correlated positively with the total score of CU traits (*r* = 0.17, *p* < .05), while associations with CBCL rule-breaking and aggression subscales, callousness, uncaring, and unemotional scores, RA and PA scores along with ADHD inattention, hyperactivity, and impulsivity scores did not reach significance or trend level. Within cases, there was no significant or trend level correlation between mean RMS-FD and these clinical characteristics reflecting severity.

In our sensitivity analyses, there was no significant influence of sex on rsFC patterns of cases compared to controls and in most of the observed subtype-specific patterns within cases (*p* > .05), aside from the RA-related right anterior insula seed-based connectivity (*p* < .001) and an influence of sex at trend-level for one uncaring-related left anterior insula seed-based connectivity (*p* = .06). Thus, we included this variable as covariate in all rsFC analyses besides age, IQ, medication, and handedness.

Thirty-eight cases had an aggression score in the clinical range but no DSM-diagnosis. These cases exhibited comparable RA scores (*M* = 12.67, *SD* = 5.62) and CU traits (*M* = 33.61, *SD* = 10.01), and lower PA scores (*M* = 3.91, *SD* = 4.00) compared to cases with a diagnosis (RA: *M* = 12.50, *SD* = 4.87; PA: *M* = 5.27, *SD* = 5.37; CU traits: *M* = 33.71, *SD* = 10.25).

We conducted sensitivity analyses for case-control group comparisons excluding the 38 cases without a DSM-diagnosis of correlations. Cases exhibited a reduced PCC seed based rsFC (*M* = 0.20) compared to controls (*M* = 0.30; *U* = 3250, *p* < .001) also present after exclusion of cases without a diagnosis (*U* = 2297, *p* < .001) and similar rsFC results for cases (*M* = 0.20) and controls (*M* = 0.30). After additional correction for ADHD scores, cases showed a diminished left anterior insula rsFC (*M* = 0.04) compared to controls (*M* = 0.10; *U* = 3739, *p* < .001), which also survived exclusion of cases without a diagnosis (*U* = 2690, *p* < .01) with comparable results for cases (*M* = 0.05) and controls (*M* = 0.10).

As cases and controls differed in average RMS-FD values, we conducted sensitivity analyses controlling for effects of this motion parameter on our case-control differences in rsFC. The PCC seed-based rsFC with a left frontal pole cluster yielded comparable statistics (*t*(196) = 5.19, cluster-size *p*-FWE < .008, peak *p*-uncorrected < .00001, *β* = .10), and the left anterior insula seed-based rsFC with a left hemispheric cluster extending from OFC to frontal pole (when additionally controlled for ADHD symptoms) survived as well (*t*(193) = 5.08, cluster-size *p*-FWE < .008, peak *p*-uncorrected < .00001, *β* = .10).

| **TABLE S3.** Distribution of demographic characteristics, diagnoses and aggression scores across sites. | | | | |
| --- | --- | --- | --- | --- |
|  |  | Cases (*n* = 118) | HC (*n* = 89) |  |
| Nijmegen (*n* = 40) | Age | 13.55 ± 2.59 | 12.64 ± 1.96 | 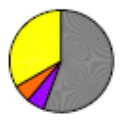 |
|  | IQ | 100.41 ± 11.83 | 107.86 ± 12.45 |  |
|  | Sex, m/f | 14/4 | 14/8 |  |
|  | Medication | 10 | 0 |  |
| Groningen  (*n* = 16) | Age | 14.77 ± 2.54 | 12.39 ± 2.32 | 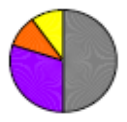 |
|  | IQ | 101.40 ± 11.63 | 101.85 ± 2.58 |  |
|  | Sex, m/f | 8/2 | 2/4 |  |
|  | Medication | 6 | 0 |  |
| Mannheim  (*n* = 33) | Age | 12.78 ± 2.42 | 12.97 ± 3.06 | 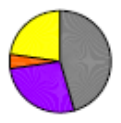 |
|  | IQ | 102.16 ± 10.13 | 116.32 ± 7.73 |  |
|  | Sex, m/f | 19/3 | 9/2 |  |
|  | Medication | 14 | 0 |  |
| Ulm (*n* = 18) | Age | 10.55 ± 2.38 | 13.59 ± 3.26 | 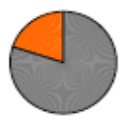 |
|  | IQ | 103.62 ± 17.55 | 101.41 ± 7.97 |  |
|  | Sex, m/f | 5/0 | 4/9 |  |
|  | Medication | 5 | 0 |  |
| London (*n* = 26) | Age | 14.67 ± 2.25 | 13.71 ± 2.05 | 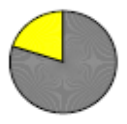 |
|  | IQ | 97.16 ± 10.33 | 111.85 ± 11.01 |  |
|  | Sex, m/f | 15/0 | 8/3 |  |
|  | Medication | 12 | 1 |  |
| Barcelona  (*n* = 23) | Age | 12.99 ± 2.82 | 14.94 ± 2.31 | 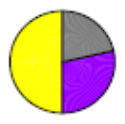 |
|  | IQ | 102.57 ± 12.10 | 105.70 ± 6.59 |  |
|  | Sex, m/f | 10/4 | 5/4 |  |
|  | Medication | 7 | 0 |  |
| Madrid (*n* = 20) | Age | 14.29 ± 2.22 | 14.68 ± 1.84 | 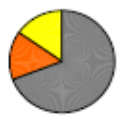 |
|  | IQ | 98.73 ± 9.45 | 105.74 ± 9.31 |  |
|  | Sex, m/f | 10/3 | 5/2 |  |
|  | Medication | 10 | 0 |  |
| Zürich  (*n* = 20) | Age | 10.53 ± 1.87 | 11.75 ± 1.49 | 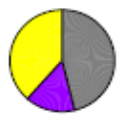 |
|  | IQ | 102.61 ± 11.20 | 99.60 ± 7.87 |  |
|  | Sex, m/f | 10/3 | 3/4 |  |
|  | Medication | 2 | 0 |  |
| Rome  (*n* = 11) | Age | 14.37 ± 2.60 | 16.12 ± 1.81 | 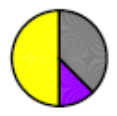 |
|  | IQ | 95.84 ± 8.83 | 97.23 ± 2.78 |  |
|  | Sex, m/f | 8/8 | 1/2 |  |
|  | Medication | 4 | 0 |  |
| Values are means ± SD, or counts. HC = healthy controls.  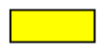 ODD  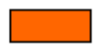 CD  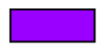 ODD + CD  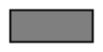 (additional) aggression in a clinical range (T > 70 on aggression subscales in Child Behavior Checklist)  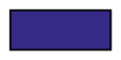Nijmegen 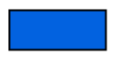 Groningen 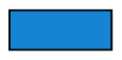Mannheim 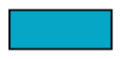 Ulm 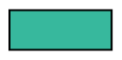 London 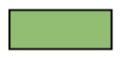 Barcelona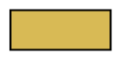 Madrid 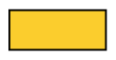 Zürich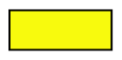 Rom  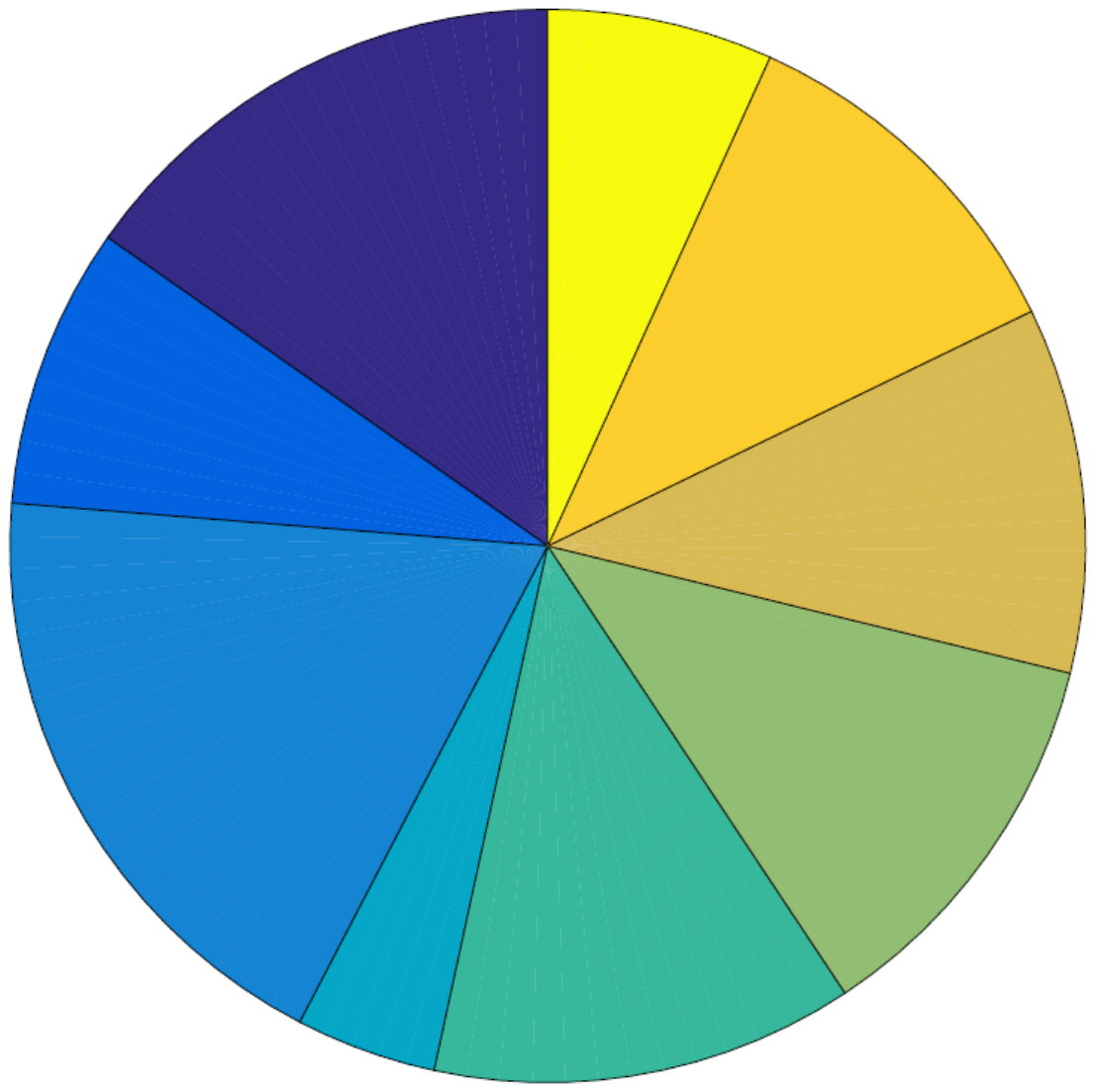  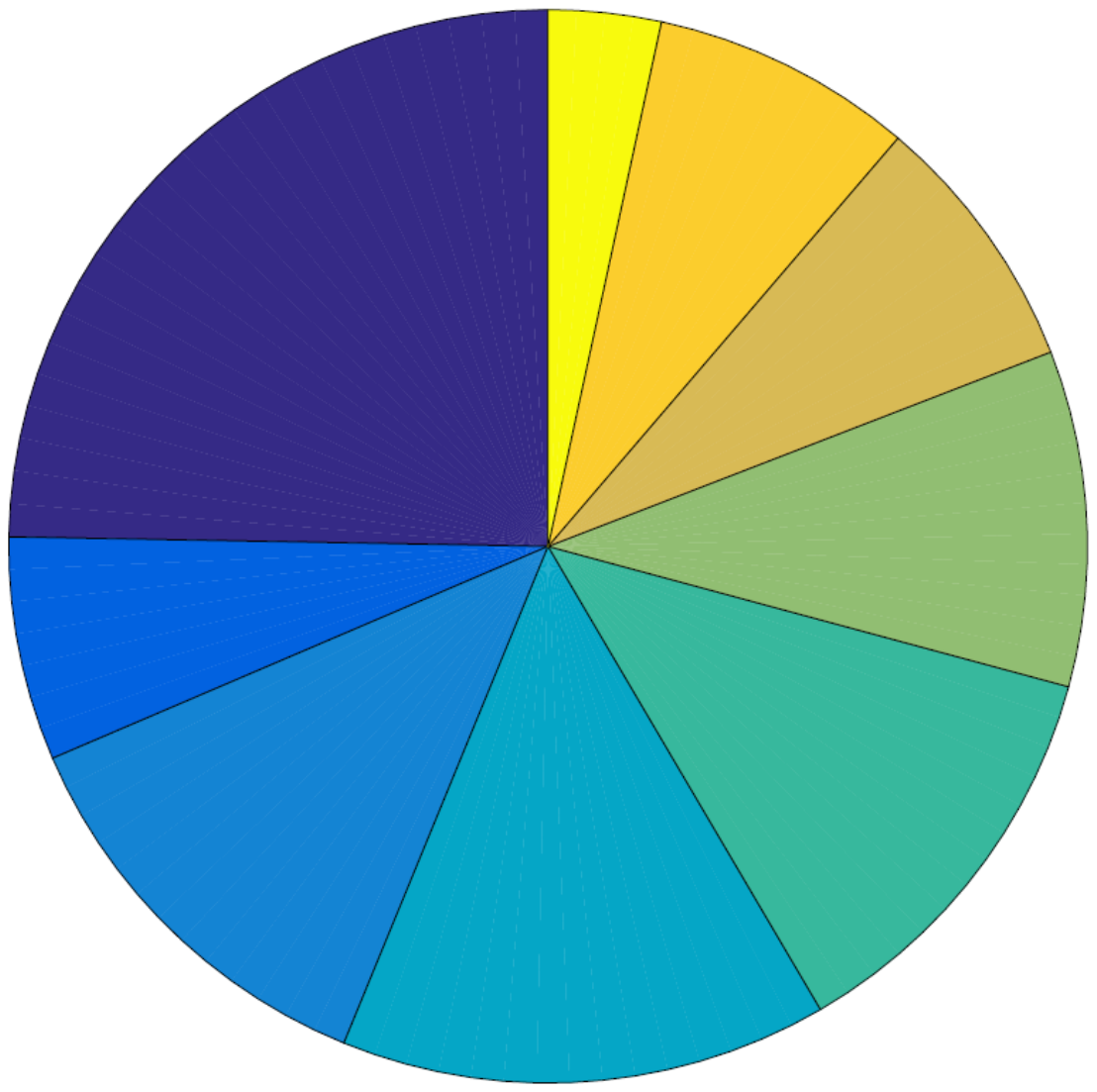 | | | | |

| **TABLE S4.** Clusters and coordinates derived from correlation analyses of seed-to-voxel connectivity in cases compared to healthy controls. | | | | | | | | |
| --- | --- | --- | --- | --- | --- | --- | --- | --- |
|  |  | | | | | Peak voxel | | |
|  |  |  |  |  |  | MNI coordinates | | |
|  | Region | Hemisphere |  | Voxels | *β*-value | x | y | z |
| 1 | *PCC Positive Association* - Frontal Pole | L | HC > cases | 225 | 0.10 | -08 | 70 | 12 |
| 2 | *Left Anterior Insula Positive Association* - Posterior Orbital Gyrus | L | HC > cases | 162 | 0.10 | -28 | 32 | -16 |
| Peak voxels are labeled according to the Anatomy toolbox. HC = Healthy Controls; PCC = Posterior Cingulate Cortex. The statistical threshold for the reported results is *p* < .001, FWE cluster-level corrected (*p* < .008 = .05/6 using additional Bonferroni correction for number of seeds) for multiple comparisons. *β*-values represent z-transformed correlation coefficients (Fisher-Z-scores). 1: Group differences corrected for site, 2: Group differences corrected for site and ADHD scores, respectively. | | | | | | | | |

| **TABLE S5.** Clusters and coordinates derived from sensitivity correlation analyses of seed-to-voxel connectivity in cases compared to healthy controls (HC) after exclusion of one site with a sample size of n < 5 for HC. | | | | | | | | |
| --- | --- | --- | --- | --- | --- | --- | --- | --- |
|  |  | | | | | Peak voxel | | |
|  |  |  |  |  |  | MNI coordinates | | |
| Region | | Hemisphere |  | Voxels | *β*-value | x | y | z |
| *PCC Positive Association* - not labeled | | L | HC > cases | 246 | 0.11 | -08 | 68 | 12 |
| Peak voxels are labeled according to the Anatomy toolbox. HC = Healthy Controls; PCC = Posterior Cingulate Cortex. The statistical threshold for the reported results is *p* < .001, FWE cluster-level corrected (*p* < .008 = .05/6 using additional Bonferroni correction for number of seeds) for multiple comparisons. *β*-values represent z-transformed correlation coefficients (Fisher-Z-scores). Group differences are corrected for site. | | | | | | | | |

| **TABLE S6.** Clusters and coordinates derived from correlation analyses of seed-to-voxel connectivity and proactive and reactive aggression scores within cases. | | | | | | | | |
| --- | --- | --- | --- | --- | --- | --- | --- | --- |
|  |  | | | | | Peak voxel | | |
|  |  |  |  |  |  | MNI coordinates | | |
|  | Region | Hemisphere |  | Voxels | *β*-value | x | y | z |
| A | *PCC Positive Association* - Calcarine Gyrus | R | Proactive Aggression | 327 | 0.07 | 04 | -66 | 18 |
|  | *Left Amygdala  Positive Association* - Precuneus  - Superior Frontal Gyrus | L R |  | 396 237 | 0.04 0.04 | -08 28 | -56 58 | 42 24 |
|  | *Right Anterior Insula  Positive Association*  - Postcentral Gyrus | L |  | 280 | 0.04 | -30 | -42 | 62 |
|  | *PCC Positive Association* - Fusiform Gyrus | R | Reactive Aggression | 249 | 0.05 | -36 | -28 | -22 |
|  | *Left Amygdala  Positive Association* - Precuneus | L |  | 864 | 0.05 | -10 | -68 | 56 |
|  | *Right Anterior Insula  Negative Association* - Calcarine Gyrus | L |  | 393 | -0.05 | -04 | -58 | 10 |
| B | *Left Amygdala Positive Association*  - Midcingulate Cortex | L | Proactive Aggression | 305 | 0.04 | -06 | -42 | 44 |
|  | *PCC Positive Association* - Fusiform Gyrus | L | Reactive Aggression | 178 | 0.05 | -28 | -20 | -28 |
|  | *Left Amygdala  Positive Association* - Precuneus | L |  | 900 | 0.05 | -10 | -68 | 56 |
|  | *Right Anterior Insula Positive Association* - not labeled | R |  | 191 | 0.05 | 24 | 06 | 28 |
| Peak voxels are labeled according to the Anatomy toolbox. PCC = Posterior Cingulate Cortex. The statistical threshold for the reported results is *p* < .001, FWE cluster-level corrected (*p* < .008 = .05/6 using additional Bonferroni correction for number of seeds) for multiple comparisons. *β*-values represent z-transformed correlation coefficients (Fisher-Z-scores). A: Main effects of proactive respectively reactive aggression corrected for site, B: Main effects of proactive respectively reactive aggression corrected for site, age, sex, IQ, medication, and handedness. | | | | | | | | |

| **TABLE S7.** Clusters and coordinates derived from correlation analyses of seed-to-voxel connectivity and callous-unemotional dimensions within cases. | | | | | | | | |
| --- | --- | --- | --- | --- | --- | --- | --- | --- |
|  |  | | | | | Peak voxel | | |
|  |  |  |  |  |  | MNI coordinates | | |
|  | Region | Hemisphere |  | Voxels | *β*-value | x | y | z |
| A | *Anterior Medial Prefrontal Cortex Positive Association* - Putamen | R | Total score | 241 | 0.05 | 26 | 16 | 00 |
|  | - Superior Parietal Lobule - Precuneus | L R | Callousness | 785 442 | 0.06 0.06 | -20 12 | -56 -66 | 66 58 |
|  | *Negative Association -* not labeled | L | Uncaring | 183 | -0.05 | -24 | -10 | 28 |
|  | *Positive Association* - Cerebelum (VIII) | R | Unemotional | 195 | 0.04 | 30 | -62 | -46 |
|  | *PCC Positive Association* - Precentral Gyrus | L | Total score | 297 | 0.07 | -54 | 02 | 48 |
|  | - pMFC - Postcentral Gyrus | L L | Callousness | 379 185 | 0.05 0.05 | -04 -36 | -06 -18 | 60 48 |
|  | *Left Amygdala Positive Association* - ACC | L | Uncaring | 171 | 0.05 | -14 | 38 | 02 |
|  | *Left Anterior Insula  Positive Association* - pMFC | L | Total score | 797 | 0.05 | -06 | -16 | 66 |
|  | - Postcentral Gyrus  - PCC - Precentral Gyrus | L L R | Uncaring | 321 285 238 | 0.06 0.07 0.06 | -20 -02 22 | -32 -42 -30 | 70 30 72 |
|  | - Precuneus - Angular Gyrus | L L | Unemotional | 544 187 | 0.05 0.05 | -06 -44 | -54 -64 | 16 32 |
|  | *Right Anterior Insula  Positive Association* - Paracentral Lobule | L | Total score | 1733 | 0.06 | -04 | -20 | 72 |
|  | - Paracentral Lobule | L | Uncaring | 370 | 0.06 | -06 | -22 | 76 |
| B | *Anterior Medial Prefrontal Cortex Positive Association*  - Inferior Parietal Lobule | L | Total score | 246 | 0.07 | -32 | -56 | 44 |
|  | - Precuneus - Superior Parietal Lobule | R L | Callousness | 1309 357 | 0.07 0.07 | 12 -28 | -66 -56 | 58 56 |
|  | - Lobule VIIb (Hem) | R | Uncaring | 188 | 0.06 | 32 | -66 | -60 |
|  | *PCC Positive Association* - Precentral Gyrus - IFG (pars orbitalis) | L L | Total score | 247 193 | 0.08 0.07 | -54 -46 | 02 12 | 48 00 |
|  | - Midcingulate Cortex | L | Callousness | 254 | 0.06 | -02 | -14 | 50 |
|  | *Left Anterior Insula  Positive Association* - pMFC | L | Total score | 858 | 0.06 | -08 | -18 | 64 |
|  | - Precuneus - Angular Gyrus | L L | Unemotional | 394 171 | 0.06 0.06 | -06 -44 | -54 -64 | 14 32 |
|  | - Precentral Gyrus - PCC | L L | Uncaring | 552 279 | 0.07 0.08 | 18 -02 | -30 -44 | 72 28 |
|  | *Right Anterior Insula Positive Association* - not labeled | L | Total score | 422 | 0.06 | -02 | -26 | 70 |
|  | - Paracentral Lobule | L | Uncaring | 183 | 0.07 | -08 | -22 | 76 |
| Peak voxels are labeled according to the Anatomy toolbox. PCC = Posterior Cingulate Cortex, IFG = Inferior Frontal Gyrus, pMFC = Posterior Medial Frontal Cortex, ACC = Anterior Cingulate Cortex. The statistical threshold for the reported results is *p* < .001, FWE cluster-level corrected (*p* < .008 = .05/6 using additional Bonferroni correction for number of seeds) for multiple comparisons. *β*-values represent z-transformed correlation coefficients (Fisher-Z-scores). A: Main effects of callous-unemotional dimensions corrected for site, B: Main effects of callous-unemotional dimensions corrected for site, age, sex, IQ, medication, and handedness. | | | | | | | | |

| **TABLE S8.** Bivariate correlations of aggression-related scores within cases. | | | | | | | | | | |
| --- | --- | --- | --- | --- | --- | --- | --- | --- | --- | --- |
|  | rule-breaking | aggres-sion | ODD | CD | in-attention | hyper-activity | impul-sivity | CU traits | reactive aggres-sion | proactive aggres-sion |
| rule-breaking |  | *r* = 0.47,  *p* < .001 | *r* = 0.16,   *p >* .05 | *r* = 0.38,  *p* < .001 | *r* = 0.12,  *p* > .05 | *r* = 0.21,  *p* > .05 | *r* = 0.04,  *p* > .05 | *r* = 0.16, *p* > .05 | *r* = 0.05, *p* > .05 | *r* = 0.14,  *p* > .05 |
| aggres-sion | *r* = 0.47, *p* < .001 |  | *r* = 0.20,  *p* < .05 | *r* = 0.21,  *p* < .05 | *r* = 0.12,  *p* > .05 | *r* = 0.09,  *p* > .05 | *r* = 0.14,  *p* > .05 | *r* = 0.34,  *p* < .001 | *r* = 0.16,  *p* > .05 | *r* = 0.22,  *p* < .05 |
| ODD | *r* = 0.16, *p >* .05 | *r* = 0.20,  *p* < .05 |  | *r* = 0.28,  *p* < .01 | *r* = 0.33,  *p* < .001 | *r* = 0.28,  *p* < .01 | *r* = 0.42,  *p* < .001 | *r* = 0.18,   *p >* .05 | *r* = 0.07,  *p* > .05 | *r* = 0.22,  *p* < .05 |
| CD | *r* = 0.47, *p* < .001 | *r* = 0.21,  *p* < .05 | *r* = 0.28,  *p* < .01 |  | *r* = 0.10,  *p* > .05 | *r* = 0.05,  *p* > .05 | *r* = 0.07,  *p* > .05 | *r* = 0.26,  *p* < .01 | *r* = 0.13,  *p* > .05 | *r* = 0.29,  *p* < .01 |
| in-attention | *r* = 0.12, *p* > .05 | *r* = 0.12,  *p* > .05 | *r* = 0.33,  *p* < .001 | *r* = 0.10,  *p* > .05 |  | *r* = 0.70,  *p* < .001 | *r* = 0.68,  *p* < .001 | *r* < 0.01,  *p* > .05 | *r* = 0.11,  *p* > .05 | *r* = -0.12,  *p* > .05 |
| hyper-activity | *r* = 0.02, *p >* .05 | *r* = 0.09,  *p* > .05 | *r* = 0.28,  *p* < .01 | *r* = 0.05,  *p* > .05 | *r* = 0.70,  *p* < .001 |  | *r* = 0.72,  *p* < .001 | *r* = -0.05,  *p* > .05 | *r* = 0.06,  *p* > .05 | *r* = 0.01,  *p* > .05 |
| impul-sivity | *r* = 0.04, *p* > .05 | *r* = 0.14,  *p* > .05 | *r* = 0.42,  *p* < .001 | *r* = 0.07,  *p* > .05 | *r* = 0.68,  *p* < .001 | *r* = 0.71,  *p* < .001 |  | *r* = -0.08,  *p* > .05 | *r* = 0.08,  *p* > .05 | *r* < 0.01,  *p* > .05 |
| CU traits | *r* = 0.16, *p* > .05 | *r* = 0.34,  *p* < .001 | *r* = 0.18,   *p >* .05 | *r* = 0.26,  *p* < .01 | *r* < 0.01,  *p* > .05 | *r* = -0.05,  *p* > .05 | *r* = -0.08,  *p* > .05 |  | *r* = 0.13,  *p* > .05 | *r* = 0.33,  *p* = .001 |
| reactive  aggres-sion | *r* = 0.05, *p* > .05 | *r* = 0.16,  *p* > .05 | *r* = 0.07,  *p* > .05 | *r* = 0.13,  *p* > .05 | *r* = 0.11,  *p* > .05 | *r* = 0.06,  *p* > .05 | *r* = 0.08,  *p* > .05 | *r* = 0.13,  *p* > .05 |  | *r* = 0.63,  *p* < .001 |
| proactive aggres-sion | *r* = 0.14, *p* > .05 | *r* = 0.22,  *p* < .05 | *r* = 0.22,  *p* < .05 | *r* = 0.29,  *p* < .01 | *r* = -0.12,  *p* > .05 | *r* = 0.01,  *p* > .05 | *r* < 0.01,  *p* > .05 | *r* = 0.33,  *p* = .001 | *r* = 0.63,  *p* < .001 |  |
| Rule-breaking and aggression T-scores are derived from the Child Behavior Checklist. ODD, CD, and ADHD hyperactivity, inattention and impulsivity scores according to the Kiddie-Schedule for Affective Disorders and Schizophrenia, present and lifetime version. Callous-unemotional (CU) traits, total score, derived from the parent-reported Inventory of Callous-Unemotional traits, reactive and proactive aggression scores according to the self-reported Reactive-Proactive Aggression Questionnaire. | | | | | | | | | | |

| **TABLE S9.** Sensitivity Analyses: Clusters and coordinates derived from correlation analyses of seed-to-voxel connectivity in cases compared to healthy controls after excluding subjects exceeding the motion threshold of >0.5 mm RMS-FD. | | | | | | | | |
| --- | --- | --- | --- | --- | --- | --- | --- | --- |
|  |  | | | | | Peak voxel | | |
|  |  |  |  |  |  | MNI coordinates | | |
|  | Region | Hemisphere |  | Voxels | *β*-value | x | y | z |
| 1 | *PCC Positive Association* - Middle Orbital Gyrus | L | HC > cases | 251 | 0.10 | -06 | 62 | 02 |
| 2 | *Left Anterior Insula Positive Association* - Inferior Frontal Gyrus, pars orbitalis | L | HC > cases | 206 | 0.11 | -30 | 34 | -14 |
| Peak voxels are labeled according to the Anatomy toolbox. HC = Healthy Controls; PCC = Posterior Cingulate Cortex. The statistical threshold for the reported results is *p* < .001, FWE cluster-level corrected (*p* < .008 = .05/6 using additional Bonferroni correction for number of seeds) for multiple comparisons. *β*-values represent z-transformed correlation coefficients (Fisher-Z-scores). 1: Group differences corrected for site, 2: Group differences corrected for site and ADHD scores, respectively. | | | | | | | | |

| **TABLE S10.** Sensitivity Analyses: Clusters and coordinates derived from correlation analyses of seed-to-voxel connectivity and reactive aggression scores within cases after excluding cases exceeding the motion threshold of >0.5 mm RMS-FD. | | | | | | | | |
| --- | --- | --- | --- | --- | --- | --- | --- | --- |
|  |  | | | | | Peak voxel | | |
|  |  |  |  |  |  | MNI coordinates | | |
|  | Region | Hemisphere |  | Voxels | *β*-value | x | y | z |
|  | *Left Amygdala Positive Association*  - Supramarginal Gyrus* - Midcingulate Cortex* - Middle Frontal Gyrus - Supramarginal Gyrus* - Middle Frontal Gyrus* | L L R R R | Proactive Aggression | 895 871 855 632 577 | 0.03 0.03 0.03 0.03 0.03 | -52 -08 34 50 28 | -26 -42 06 -40 62 | 26 44 48 44 22 |
|  | *PCC Positive Association* - Fusiform Gyrus* | L | Reactive Aggression | 1016 | 0.05 | -28 | -22 | -28 |
|  | *Left Amygdala  Positive Association* - Precuneus - Angular Gyrus - not labeled - Supramarginal Gyrus | L R R L |  | 1028 300 242 208 | 0.05 0.05 0.05 0.05 | -06 32 34 -56 | -70 -62 04 -28 | 60 54 46 42 |
|  | *Right Amygdala Positive Association* - Superior Frontal Gyrus | L |  | 156 | 0.05 | -18 | 58 | 24 |
| Peak voxels are labeled according to the Anatomy toolbox. PCC = Posterior Cingulate Cortex. The statistical threshold for the reported results is *p* < .001 and for * p < .01, FWE cluster-level corrected (*p* < .008 = .05/6 using additional Bonferroni correction for number of seeds) for multiple comparisons. *β*-values represent z-transformed correlation coefficients (Fisher-Z-scores). Main effects of reactive aggression corrected for site, age, sex, IQ, medication, and handedness. | | | | | | | | |

| **TABLE S11.** Sensitivity Analyses: Clusters and coordinates derived from correlation analyses of seed-to-voxel connectivity and callous-unemotional dimensions within cases after excluding cases exceeding the motion threshold of >0.5 mm RMS-FD. | | | | | | | | |
| --- | --- | --- | --- | --- | --- | --- | --- | --- |
|  |  | | | | | Peak voxel | | |
|  |  |  |  |  |  | MNI coordinates | | |
|  | Region | Hemisphere |  | Voxels | *β*-value | x | y | z |
|  | *Anterior Medial Prefrontal Cortex Positive Association*  - Superior Orbital Gyrus* | L | Total score | 538 | 0.06 | -20 | 30 | -14 |
|  | - Postcentral Gyrus - Precuneus | R R | Callousness | 358 170 | 0.08 0.07 | 32 08 | -40 -68 | 46 56 |
|  | - not labeled | R | Uncaring | 218 | 0.06 | 32 | -68 | -60 |
|  | *PCC Positive Association* - Pallidum | L | Total score | 305 | 0.06 | -20 | -06 | 06 |
|  | - Middle Temporal Gyrus* | L | Callousness | 805 | 0.06 | -52 | -42 | 06 |
|  | - Inferior Frontal Gyrus, pars triangularis | L | Uncaring | 169 | 0.09 | -34 | 32 | 12 |
|  | *Left Anterior Insula  Positive Association* - Precentral Gyrus - PCC | R L | Uncaring | 530 286 | 0.07 0.09 | 22 -02 | -30 -46 | 72 34 |
|  | - Paracentral Lobule - Precuneus | L L | Unemotional | 440 221 | 0.05 0.06 | -10 -06 | -32 -54 | 68 16 |
|  | *Right Anterior Insula Positive Association* - Paracentral Lobe | L | Total score | 221 | 0.06 | 00 | -24 | 70 |
|  | - Rectal Gyrus* | L | Uncaring | 598 | 0.06 | -10 | 24 | -12 |
| Peak voxels are labeled according to the Anatomy toolbox. PCC = Posterior Cingulate Cortex. The statistical threshold for the reported results is *p* < .001 and for * p < .01, FWE cluster-level corrected (*p* < .008 = .05/6 using additional Bonferroni correction for number of seeds) for multiple comparisons. *β*-values represent z-transformed correlation coefficients (Fisher-Z-scores). Main effects of callous-unemotional dimensions corrected for site, age, sex, IQ, medication, and handedness. | | | | | | | | |
